# Early-Life Behavior Phenotypes and Cortisol Responses to Common Lab Stressors in a Cichlid Fish

**DOI:** 10.1101/2024.10.07.617115

**Authors:** Alyssa P. Alvey, Eliyah R. Stern, June Lee, Abigail G. Parrish, Tessa K. Solomon-Lane

**Author notes:** Corresponding author Dr. Tessa Solomon-Lane Department of Natural Sciences / W.M. Keck Science Department Scripps, Pitzer, and Claremont McKenna Colleges 925 N. Mills Ave, Claremont, CA 91711, USA.

## Abstract

The stress response is highly conserved across species, and increased glucocorticoid release (cortisol in fishes) is a key element. In the highly social cichlid fish, Burton’s Mouthbrooder (*Astatotilapia burtoni*), stress axis activity is associated with juvenile social behavior and status, and it mediates early-life social effects, yet little is known about early-life stress physiology. We measured water-borne cortisol, a non-invasive method, in juveniles less than 1-week old. We first tested whether juveniles habituate to the beaker confinement necessary for sample collection. Repeated exposure to a beaker did not affect cortisol compared to handled and unhandled controls. In a separate cohort, we next measured behavior in an open field exploration and social cue investigation, followed by a test of whether common lab stressors elevated cortisol. Controls were undisturbed in a collection beaker (90 min). For the stressor treatments, we collected three sequential samples (30 min each) to try and quantify stress response peak and recovery; however, we found no effect of time. Stressors included handling, brief net confinement, and brief gentle movement. These stressors led to a significant increase in cortisol, with the highest levels resulting from confinement. The behavior tests revealed a bold-shy axis across both tests, describing the behavior of most individuals. A distinct group of socially-motivated juveniles showed a dramatic switch, staying in the territory during the open field but leaving to investigate the social cue. We found no associations between behavior and cortisol response. This work provides insight into early-life behavior and stress axis development and function.

**Highlights:**

- *Astatotilapia burtoni* under 1-week old did not habituate to beaker confinement
- Young juveniles displayed bold/shy or socially-motivated behavior phenotypes
- Cortisol increased in response to lab stressors: handling, confinement, and movement
- Individual variation in cortisol establishes an expected hormone range for this age
- Juvenile cortisol did not correlate with behavior or differ by behavior phenotype

## 1. Introduction

Early-life is characterized by rapid growth and development (Forbes and Dahl, 2010), during which, experiences such as stress can have profound impacts on ontogeny (Antunes and Taborsky, 2020; Bannier et al., 2017; Baxter, 1978; Ruys et al., 2017). Life history transitions (e.g., from neonatal or larval to juvenile stages, metamorphosis, puberty) are often mediated by hormone signaling, including the neuroendocrine stress axis (Antunes et al., 2021; Antunes and Taborsky, 2020; Denver, 1997; Glennemeier and Denver, 2002; Jonsson and Jonsson, 2014; Solomon-Lane et al., 2013). These transitions are times of heightened sensitivity, when organisms undergo important biological changes at multiple levels of organization (Denver, 1997; Glennemeier and Denver, 2002). The stress response is highly evolutionarily conserved and acts on multiple timescales, enabling organisms to survive potentially dangerous or threatening situations by regulating behavior and physiology (Beery and Kaufer, 2015; Taborsky et al., 2021; Wendelaar Bonga, 1997). A stressor is a stimulus that causes (or threatens to cause) the disruption of homeostasis and examples include extreme weather changes, predation, excitement, social stress, injury, or disease (Nelson and Kriegsfeld, 2022; Taborsky et al., 2021). While the impacts of early-life stress have been extensively researched and demonstrate powerful effects on an organism’s ability to adjust to its environment (Branchi et al., 2013; Glennemeier and Denver, 2002), our understanding of stress physiology in very young organisms remains limited for many species.

Here, we focus on the hypothalamic-pituitary-adrenal (interrenal in fish and amphibians; HPA/I) axis (Nelson and Kriegsfeld, 2022; Spencer and Deak, 2017; Wendelaar Bonga, 1997). When an organism encounters a stressor, the HPA/I axis is activated, initiating the release of corticotropin-releasing factor (CRF) in the hypothalamus. This signals for the release of adrenocorticotropic hormone (ACTH) from the pituitary gland, which prompts the release of glucocorticoids (cortisol in fishes) (Spencer and Deak, 2017; Taborsky et al., 2022; Wendelaar Bonga, 1997). Cortisol aids in mobilizing energy away from processes unnecessary for immediate survival, such as digestion and growth, and towards fighting or escaping a threat (Glennemeier and Denver, 2002; Spencer and Deak, 2017). During a stress response, heart rate, blood pressure, oxygen intake, and blood glucose levels all increase; blood is directed to the muscles and away from other organs; and immune function can be affected (Glennemeier and Denver, 2002; Khulan and Drake, 2012; Nelson and Kriegsfeld, 2022; Wendelaar Bonga, 1997). Although cortisol is one of many molecules involved in the stress response, it is commonly used as a biomarker for stress (Breuner et al., 1998). Changes in cortisol levels can be used to indicate the presence, duration, and perceived intensity of a stressor (Beery et al., 2020; Dhabhar and Mcewen, 1997; Nelson and Kriegsfeld, 2022).

Characterizing the stress response during formative stages of life provides insight into HPA/I axis development and how individuals interpret and respond to their experiences. For example, some species exhibit a stress hyporesponsive period (SHRP) (M. Vázquez, 1998). During the SHRP, animals typically display low baseline cortisol levels (Butte et al., 1973) and minimal or no stress reactivity (M. Vázquez, 1998; Manolova and Gulyaeva, 2023). For example, rainbow trout (*Oncorhynchus mykiss*) experience a two week SHRP immediately after hatching and have reduced reactivity to stressors until they are about 3-4 weeks old (Barry et al., 1995). Not all species undergo the SHRP; in those that do not, there is often high individual variation in HPA/I axis function early in life (Baugh et al., 2014). This individual variation can be linked to a combination of intrinsic factors, such as behavior, as well as extrinsic environmental conditions (Cockrem, 2013). In the Japanese Meagre fish (*Argyrosomus japonicus*), boldness scores were negatively correlated with cortisol levels, and this hormone-behavior interaction was maintained even though individual phenotype varied across trials (Raoult et al., 2012). Resource availability is also relevant, and juvenile rainbow trout with limited access to food exhibited higher cortisol levels (Olsen et al., 2005). Additionally, early-life social experience also had an impact, where factors such as social stability and the presence of parental figures influenced cortisol levels (Antunes et al., 2021; Bender et al., 2008; Harmon et al., 2024; Mileva et al., 2009).

We studied Burton’s Mouthbrooder (*Astatotilapia burtoni*), a highly social cichlid fish and model system in social neuroscience, to investigate early-life stress physiology and behavior. Both adults and juveniles of this species live in social groups throughout all post-larval life history stages and express a highly conserved suite of social behaviors, such as affiliation, aggression, and territoriality (Fernald, 1977; Fernald and Hirata, 1979; Maruska and Fernald, 2018). Adults form naturalistic mixed-sex, hierarchical communities with females and males of different status classes. Juveniles readily interact socially and form status relationships in all-juvenile groups (Solomon-Lane et al., 2022). HPI axis function is important across *A. burtoni* life history stages, including for juvenile (Solomon-Lane et al., 2022) and adult social status and behavior (Alcazar et al., 2016; Carpenter et al., 2014; Dijkstra et al., 2012; Korzan et al., 2014; Maruska et al., 2022). For example, dominance in juveniles is associated with increased whole brain gene expression of glucocorticoid receptor (GR) 1a and GR1b, and decreased expression of GR2 and mineralocorticoid receptor (Solomon-Lane et al., 2022). In adults, cortisol tends to be higher in subordinate males, but not in all environments (Maruska et al., 2022). Cortisol, social behavior, and status are associated in context-specific ways (Alcazar et al., 2016; Clement et al., 2005), and neural expression of HPI axis-related genes shift in response to changes in social context (Korzan et al., 2014).

In this study, we investigated juvenile behavior and cortisol response in fish less than 1-week old. We previously showed the HPI axis to be sensitive to social effects in fish at this early age. Juvenile fish reared in social groups compared to those reared in pairs had significantly higher whole brain gene expression of GR1a at 1-week and 5-weeks old. Expression of CRF and GR2 both changed over developmental time, with significantly higher expression at 5-weeks (Solomon-Lane and Hofmann, 2019). In a separate study, social stability had a significant effect on an integrated behavior-hormone phenotype measured in 5-week-old juveniles, with higher cortisol being associated with lower social status, less bold or exploratory behavior, and more time in a territory, away from a novel social cue fish (Harmon et al., 2024). Beyond these findings, our understanding of HPI axis function in very young *A. burtoni* remains limited.

Here, we first tested how juveniles less than 1-week old respond to the water-borne hormone collection method. In adult daffodil cichlids (*Neolamprologus pulcher*) and convict cichlids (*Amatitlania nigrofasciata*), fish placed in a hormone collection beaker respond with an elevation in cortisol. However, with repeated exposure for 3-4 consecutive days, cortisol levels decreased significantly, showing that fish habituated to the experience. Thus, cortisol levels following habituation are a better indicator of baseline levels (Antunes et al., 2021; Wong et al., 2008). For our study, we exposed juvenile *A. burtoni* to a collection beaker on 3 consecutive days, starting the day they were removed from their mother’s buccal cavity. We collected cortisol samples on the fourth day and compared levels to handled and unhandled control fish. We expect all treatment groups to have detectable levels of water-borne cortisol, with habituated fish exhibiting the lowest levels.

In a separate cohort of unhabituated fish, we then tested the hypotheses that juveniles less than 1-week old would 1) show individual variation in behavior; 2) respond to common lab stressors with an increase in cortisol, and 3) variation in behavior would be associated with variation in cortisol. First, behavior was observed in an open field exploration and social cue investigation, two tests that have been used successfully with older *A. burtoni* juveniles (Harmon et al., 2024; Solomon-Lane and Hofmann, 2019) and across diverse species. The open field exploration can be used to measure activity, exploration-avoidance behavior (Conrad et al., 2011); bold-shy behavior (Gebauer et al., 2023; Yuen et al., 2017), and anxiety-like behaviors (e.g., (Cachat et al., 2010; Prut and Belzung, 2003). The social cue investigation can be used to assess social motivation or preference (e.g., (Bonuti and Morato, 2018; Moy et al., 2004)).

Immediately after the behavior tests, we exposed juveniles to a stressor and measured cortisol. The stressors included handling (“beaker”), confinement (“net”), and water movement (“orbital,” via orbital shaker). We chose these treatments because similar stimuli are used in stress research across a variety of species (Cockrem, 2013) and they are common experiences for laboratory fish. Previous studies on the habituation of fish to hormone collection beakers demonstrate that the experience can elevate cortisol (Antunes et al., 2021; Wong et al., 2008). Brief confinement has been shown to increase cortisol in many species (Ding et al., 2021; Eleftheriou et al., 2020; McGivern et al., 2009), and for fish, confinement usually takes place in a net (in or out of water) (Cerqueira et al., 2021; Jentoft et al., 2002; Ramsay et al., 2009). While using an orbital shaker to generate water movement as a stressor (Adamante et al., 2008) is not as common as beaker or net confinement, movement has been used as a stressor (Villarroel et al., 2021) and is a relevant (and unavoidable) component of handling. In the wild, water movement is natural and varies considerably across bodies of water, season, and more (**cite)**. Overall, this work has the potential to establish baseline and stress-induced ranges of cortisol in very young juveniles, creating a foundation for future research. Additionally, this information can be applied to improve lab procedures and fish welfare.

## 2. Methods

### 2.1 Study Organism

*A. burtoni* juveniles came from a lab population (see Hofmann / Renn lab strain: (Pauquet et al., 2018) that were the descendants of a wild-caught colony ∼65-70 generations prior (Fernald and Hirata, 1977). The juveniles’ parents were selected due to reproductive timing with the start of the experiments, not specifically for other attributes (e.g. age, size). The parents were housed in mixed-sex communities with other adults of similar developmental stage and size. During reproduction, the female lays her eggs in a dominant male’s territory. She then scoops them into her mouth, where the male fertilizes them. The mother incubates them orally for 10-13 days. In their natural habitat, the female would release the juveniles on her own, allowing them to swim in and out of her mouth for protection for a few more weeks. In this laboratory setting, the females do not release them, so for these experiments, we stripped the juveniles by hand ∼12 days post-fertilization, once they no longer had visible yolk-sacs and consistently swam upright (Fernald and Hirata, 1977; Fraley and Fernald, 1982). All the juveniles in these studies were under 1 week of age and came from multiple clutches across multiple aquaria. When considering age, we defined 0-days old (“d0”) as the day juveniles were removed from the mother’s buccal cavity (11-13 days post-fertilization). We are unable to reliably determine the sex until maturation; however, *A. burtoni* hatch at about a 1:1 female to male ratio (Heule et al., 2014). The juveniles were fed once daily with Hikari Middle Larval Stage Plankton (Pentair Aquatic Eco-systems). All aquaria within the laboratory were maintained at an ambient temperature of 23.3-24.4 °C and experienced a 12:12 light/dark cycle. After the fish underwent behavioral testing and/or hormone collection, the individuals were placed in separate community aquaria to avoid including any fish in these studies twice.

### 2.2 : Habituation to water-borne hormone collection methods

#### 2.2.1 Experimental juveniles and housing

A total of 39 juveniles were used to test whether fish less than 1 week old habituate to the water-borne hormone collection method (see 2.2.3). Experimental day 1 started with d0 juveniles, the day they were removed from the mother’s buccal cavity. The juveniles were housed in pairs in small acrylic aquaria (20.3 x 12.7 x 13.4 cm, 3.4 L) and assigned to a treatment group: control (n=6 pairs), handled (n=7 pairs), or habituated (n=7 pairs). The aquaria contained one small terracotta pot at one end of the aquarium. White dividers were placed between the aquaria to prevent visual contact between pairs and to reduce environmental disturbances.

#### 2.2.2 Animal Handling and Habituation

There were three treatment groups consisting of habituated (n=14), handled (n=13), and control groups (n=12) (see *2.2.3*, Figure 1A). Both fish in each pair were assigned to the same treatment group. The experimental procedure lasted four days. For the habituated group, the procedure comprised the process of water-borne hormone collection that fish in all groups experienced on the fourth day. This allowed for the repeated experience of the collection conditions, which could potentially affect hormone levels. On experimental days 1-3, the juveniles were caught with a small net (3.6 cm diameter) and transferred individually to a 20 mL beaker containing 15 mL of water, where they remained for 30 min. Beakers were placed in a cardboard grid to reduce visual disturbances. After 30 min, the fish were returned to their aquaria. For the handled group on experimental days 1-3, both fish from the aquarium were individually transferred to the same beaker of 250 mL aquarium water using a small, round net. Immediately afterwards, they were gently poured back into their aquarium. On experimental days 1-3, the control group fish remained in their aquaria with no disturbances. After hormones were collected on Day 4, we collected body mass (g).

**Figure 1:**
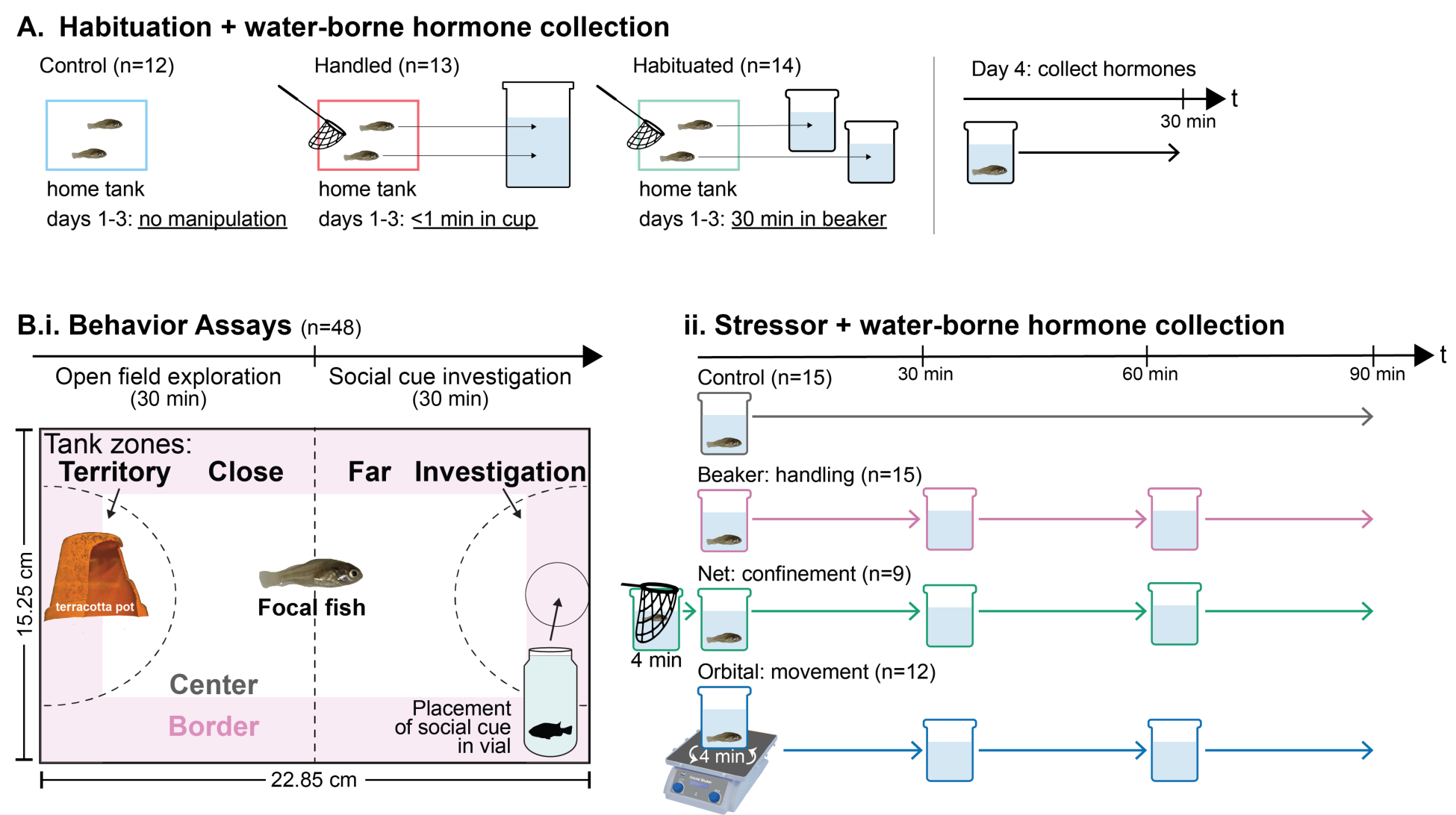
Experimental designs. **A.** Testing whether juveniles habituated to the water-borne hormone collection protocol started the day juveniles were removed from the mother’s buccal cavity (day 1). Control juveniles (n=12) were left undisturbed in their home aquarium. Handled juveniles (n=13) on days 1-3 were transferred to a 200 mL plastic beaker for > 30 s before being returned to their home aquarium. Habituated juveniles (n=14) were transferred to hormone collection beakers for 30 min on days 1-3, then returned to their home aquarium. Water-borne hormones were collected on day 4 for all fish. **B.i.** Using a separate cohort of juveniles less than 1 week old, behavior was tested right before stressor exposure. In the open field exploration (min 0-30), the focal fish was alone in the aquarium. In the social cue investigation (min 30-60), a small juvenile cue fish inside of a scintillation vial was placed in the circle in the investigation zone. The experimental aquarium contained a terracotta pot shelter/territory. The “territory,” “close,” “far,” and “investigation” zones were drawn on the aquarium. The border vs. center zones were not physically indicated. **B.ii.** Immediately after the behavior tests, juveniles were exposed to one stressor treatment for water-borne hormone collection: control (n=15), beaker (handling, n=12), net (confinement, n=9), or orbital shaker (movement, n=12).

#### 2.2.3 Hormone collection

On experimental day 4, water-borne hormones were collected from all fish in all groups. Water-borne hormones collection is a non-invasive method during which fish are placed individually in a beaker of water for an extended period of time to allow for a natural accumulation of steroid hormones excreted through the gills, urine, and feces in the water (Friesen et al., 2012; Kidd et al., 2010). We collected each cortisol sample in 15 mL of clean aquarium water in a 20 mL beaker for 30 min. Beakers were placed inside a cardboard grid to reduce visual disturbances from the side and prevent fish from seeing one another. After 30 min, fish were removed from the beakers and placed in a community aquarium. All hormones samples were collected between 1230 hrs and 1330 hrs to control for diurnal cortisol fluctuations (Gabor and Contreras, 2012). Water samples were frozen in conical tubes at -20° C until analysis (Koakoski et al., 2012; Zuberi et al., 2014).

#### 2.2.4 Processing hormone samples and measuring cortisol

Solid-phase extraction was used to isolate cortisol from the water sample using 3 cc Sep-Pak Vac C18 columns (Water Associates, Milford, MA) on a vacuum manifold. Briefly, the columns were prepared for the extraction using 2 consecutive runs of 2 mL methanol, ensuring that the columns did not dry out. Next, 2 mL of ultrapure water were passed through, twice. Then, the samples were run through the columns using the vacuum pump on the manifold. Afterwards, two more applications of 2 mL ultrapure water was passed through the columns, followed by 5 min of vacuuming to allow the columns to dry. To elute the cortisol samples into 13 x 100 mm borosilicate test tubes, 2 mL of methanol were passed through the columns, twice. The sample in methanol was frozen at -20°C until it was dried under a low-pressure stream of nitrogen at 37°C to evaporate the methanol. The samples were then resuspended to 300 μL, with 5% EtOH and 95% ELISA buffer. The samples were then further diluted 1:8, with 50 μL of resuspended sample into 400 μL ELISA buffer. The samples were shaken for 1 hr on a multitube vortex and then stored at -20°C. In preparation of the ELISA, the samples were thawed and shaken again on the multitube vortex for 1 hr. We performed the ELISA, strictly adhering to the protocol provided by the manufacturer (Cayman Chemical, Ann Arbor, Michigan). One sample was used as an interplate control. The standard curves had r^2^ values of 0.98 and 0.99. We present results as pg of cortisol per mL collection water volume per hr sample collection.

### 2.3 Testing behavior and cortisol responses to common lab stressors

#### 2.3.1 Experimental juveniles and housing

A total of 53 juveniles were removed from the mothers’ buccal cavity and immediately transferred to small acrylic community aquaria (20.3 x 12.7 x 13.4 cm, 3.4 L). Each aquarium contained a single air stone, 2 clay terracotta territories/shelters, and no substrate. Fish remained in the community aquaria until participating in the behavioral test (see *2.3.2*) and stress treatments (see *2.3.3*).

#### 2.3.2 Behavioral Tests

To quantify attributes of juvenile behavioral phenotype, we employed two behavioral tests: an open field exploration test and a social cue investigation test. The open field test has been previously used to detect and measure locomotion, exploration, and anxiety-like behaviors. Across species, higher locomotion has been correlated with higher cortisol levels, while increased time in the center of the aquarium / arena is correlated with lower cortisol levels. The social cue test has been previously used to assess social motivation and preference, quantifying sociality and territoriality. Typically, higher social motivation correlates with lower cortisol levels (Bonuti and Morato, 2018; Cachat et al., 2010; Moy et al., 2004; Prut and Belzung, 2003). We selected these two tests due to their well-established use in studying the behavior of various species, allowing for cross-species comparisons. They have also been successfully used with older (∼2 mo old) juvenile *A. burtoni* (Harmon et al., 2024; Solomon-Lane and Hofmann, 2019) compared to fish in this study.

Juvenile fish were haphazardly selected from the home aquaria, all under 7 days old. Fish that appeared visually unhealthy were not included. The individuals were placed alone into an experimental test aquarium (20.3 x 12.7 x 13.4 cm; 3.4 L) with no lid. The aquarium had been divided into four zones using black permanent marker. The territory zone, which was on one short side of the aquarium, contained a small terracotta pot territory/shelter (5 cm diameter). The investigation zone, which was on the opposite short side of the aquarium, marked the placement of the round vial containing the social cue during the social cue investigation. The zone also included a 2.54 cm area around the vial. The close zone was between the territory zone and the halfway line of the aquarium, and the far zone was between the halfway line and the investigation zone. The center and the border were scored, but not physically marked on the aquarium. The border was 2.54 cm around the edge of the aquarium. The center was defined as the rest of the aquarium (Figure 1B.i.).

Both of these tests have previously been described in detail (Solomon-Lane and Hofmann, 2019). Briefly, experimental testing lasted 1 hr. Fish were visually recorded from above the aquarium (Warrior 4.0, Security Camera Warehouse, Asheville, NC). Once a focal fish was placed in an experimental aquarium, the open field exploration began. The focal fish remained alone for the duration of the test (0-30 min). After the 30 min had elapsed, the same free-swimming focal fish remained, but a second social-cue fish was placed in the investigation zone in a scintillation vial, initiating the social cue investigation (30-60 min). Behavioral Observation Research Interactive Software (BORIS) was used to score the behavior videos (Friard and Gamba, 2016). For each test, 10 min of behavior footage was analyzed (open field: min 20-30; social cue: min 32-42). The frequency entering each zone of the aquarium and time (s) spent in each zone was scored. During the social cue investigation, only behavior from the free-swimming focal fish was measured.

#### 2.3.3 Stress Treatments and Water-Borne Cortisol Collection

Immediately after the behavior tests, each individual fish was assigned to a stressor treatment (see *2.3.4*) and water-borne steroid hormones were collected between 1100 to 1330 hrs (Gabor and Contreras, 2012). During each cortisol sample collection, the collection beaker was set inside a white paper barrier to reduce disturbance from visual stimuli. Following collection (see below), water samples were frozen at -20° C until further analysis.

##### 2.3.3.1 Control

In the control treatment, individual fish were placed in 15 mL of clean aquarium water in 20 mL beakers and left undisturbed for 90 min. To collect water-borne hormones, a minimum amount of handling must occur. The control group is the most minimally manipulated treatment group, while still meeting the requirements for hormone collection. Each stressor treatment (see below: beaker, net, orbital) builds on the control treatment by adding specific, additional stimuli to the fish. We did not habituate the fish to this protocol because juveniles this young did not have significantly lower cortisol following habituation (see Results *3.1*).

##### 2.3.3.2 Beaker Treatment

To measure the impact of the additional handling that is necessary for collecting sequential time-points of water-borne hormones, individual fish were placed in 15 mL of clean aquarium water in a 20 mL collection beaker and then transferred into a fresh beaker of aquarium water every 30 min for a total of 90 min. Water-borne samples were collected from each of the 3 time points.

##### 2.3.3.3 Net Treatment

To determine the effects of confinement stress within a fish net on cortisol levels, juveniles were placed individually in a fish net (12.7 x 15.2 cm frame), such that the net reached to the bottom of a 140 mL beaker with 120 mL of water. They were kept in the net for four min, then transferred to the first hormone collection beaker (15 mL of clean aquarium water in a 20 mL beaker) for 30 min. The second and third water samples collections were the same as for the beaker treatment.

##### 2.3.3.4 Orbital Treatment

To measure the impact of small-scale water turbulence on the cortisol levels of juveniles, each individual was placed in their first hormone collection beaker on top of an orbital shaker (Corning LSE Low Speed Orbital Shaker) rotating slowly at 30 rpm for 4 min. The shaker was then turned off, and the fish spent the remaining 26 min in the same collection beaker. The second and third cortisol sample collections were the same as for the beaker treatment.

On the first and second rounds of our experiment, we used a Penn-Plax Quick Net fish net (12.7×15.2 cm) to transport the fish between hormone collection beakers. For the third round, we used custom-made, spoon-shaped hand nets that were small (3.6 cm diameter) and shallow. Nets of this size and shape were safer for fish of this small size and prevented them getting stuck in the corners.

#### 2.3.4 Processing hormone samples and measuring cortisol

Water-borne hormones were extracted and quantified using ELISA as in *2.2.4*. The samples were then resuspended to 300 μL, with 5% EtOH and 95% ELISA buffer. The samples were then further diluted 1:8, with 50 μL of resuspended sample into 400 μL ELISA buffer. We used two samples as an interplate control across the four ELISA plates. Values on one plate were substantially higher than the three plates; therefore, we used a scaling factor to equalize the plates to one of the interplate controls.

### 2.4 Statistical Analysis

All statistical analyses were performed using the R software version 4.4.0 (RStudio Team, 2022). Results were considered significant at the *p*<0.05 level, and averages ± standard error of the mean are included in the text. The boxes of the box and whisker plots show the median and the first and third quartiles. The whiskers extend to the largest and smallest observations, within or equal to 1.5 times the interquartile range. To test for differences in cortisol levels based on treatment or behavior type, we used one-way ANOVA for data that met the assumptions of parametric statistics. A Kruskal-Wallis test was used for data that did not meet the assumptions of parametric statistics. Dunn’s tests were used for *post hoc* analysis of significant ANOVA results, and effect size was measured using eta-squared (small effect: 0 < η2 < 0.01; moderate: 0.01 < η2 < 0.06; large: 0.06 < η2). We used a two-way mixed effects ANOVA to test for the effects of treatment (independent measure) and collection time period (repeated measure) on cortisol levels. Differences in the time and frequency in each aquarium zone in the open field exploration and social cue investigation were analyzed using Friedman tests, with *post hoc* analysis of significant results using Wilcoxon tests with Bonferroni correction for multiple comparisons. Kendall’s W is reported for effect size (small effect: 0.1 - < 0.3; moderate: 0.3 - < 0.5; large: ≥ 0.5). To compare between the border and center zones, and between the open field and social cue tests, we used Wilcoxon signed-rank tests and measured effect size ‘r’ (small effect: 0.1 - < 0.3; moderate: 0.3 - < 0.5; large: ≥ 0.5). Independent t-tests were used to test whether net size or experiencing a fall during hormone collection affected cortisol levels. Pearson correlations were used to test whether behavior variables, including locomotion, time in the center of the aquarium during the open field or social cue tests, or PCs (see below), were significantly associated with cortisol. Finally, we used Principal Components Analysis (PCA) to identify how behavior in the open field exploration and social cue investigation clustered, including time in the territory, close, far, investigation, and center zones. We also split PC2 into quartiles, and we used Pearson correlations to test for associations among times in different zones of the aquarium during the open field exploration and social cue investigation. Data from fish that did not survive through all phases of the experiment were not included in the analysis.

## 3. Results

### 3.1 Effects of habituation on water-borne cortisol

We first compared cortisol levels in control, handled, and habituated juveniles. We found no significant differences in cortisol across treatment groups (F(2,36)=1.11, p=0.90, Fig 2).

**Figure 2:**
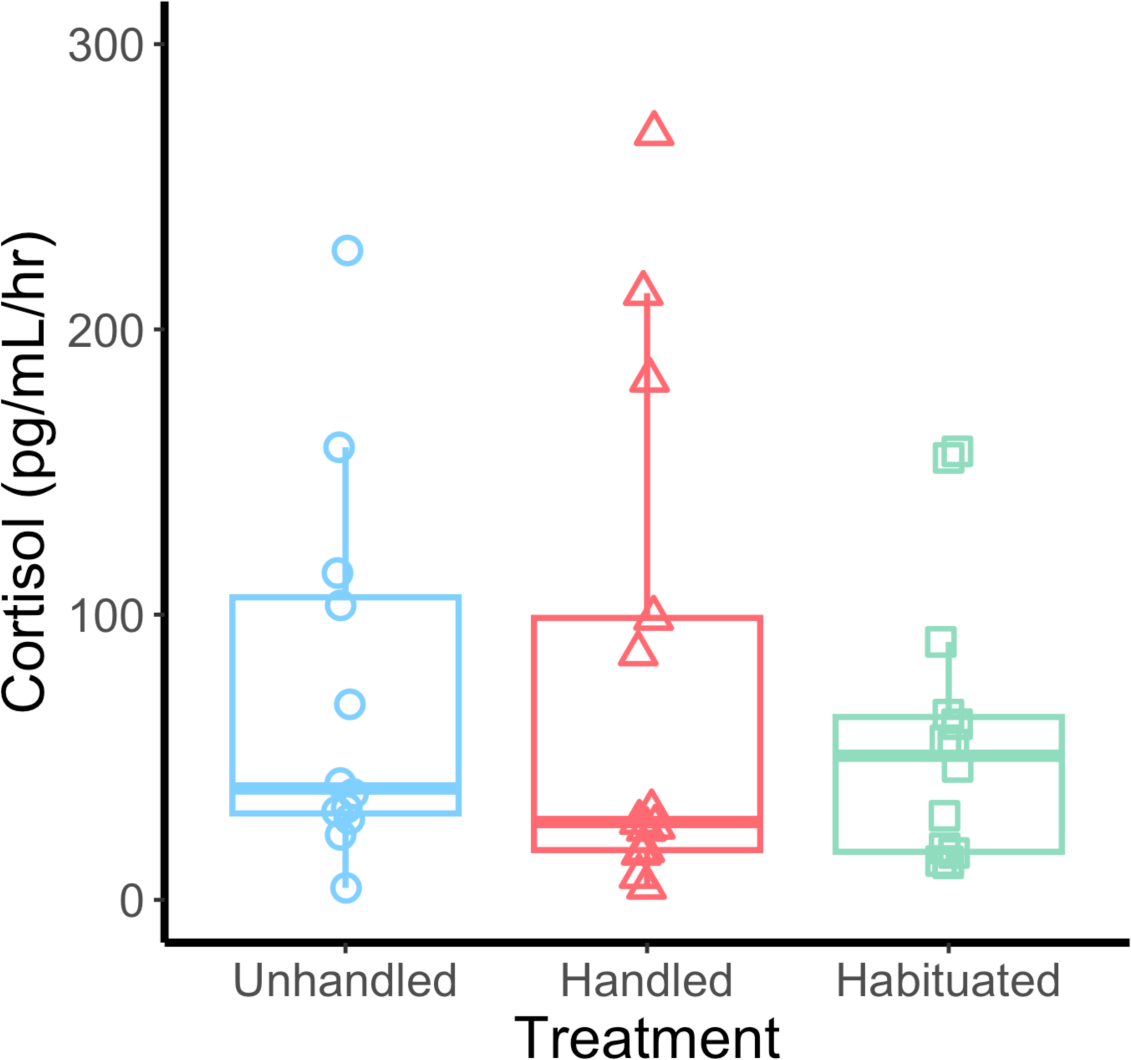
Water-borne cortisol levels (pg/mL/hr) in unhandled control (n=12), handled control (n=13), and habituated (n=14) juveniles.

### 3.2 Open field exploration and social cue investigation behavior

In a separate cohort of juveniles, we observed behavior in open field exploration and social cue investigation tests, after which, fish were exposed to lab stressors and water-borne hormone samples were collected. In both the open field exploration (χ²(3)=47.0, p<0.0001, W=0.33) and social cue investigation (χ²(3)=33.4, p<0.0001, W=0.23), individuals spent significantly different amounts of time in the different zones of the aquarium (Fig 3). In the open field exploration, juveniles spent the most time in the territory zone and the least time in the investigation zone. In the social cue investigation, fry spent the most time in the investigation zone and the least time in the territory zone. Table 1 shows the results of *post hoc* pairwise comparisons between zones for both behavior tests. Duration in the center versus the border was analyzed separately because the center and border overlap with the other aquarium zones (territory, close, far, and investigation) (Fig 1A). In both the open field exploration (p<0.0001, r =0.57) and social cue investigation (p=0.013, r=0.36), juveniles spent significantly more time in the border zone.

**Figure 3:**
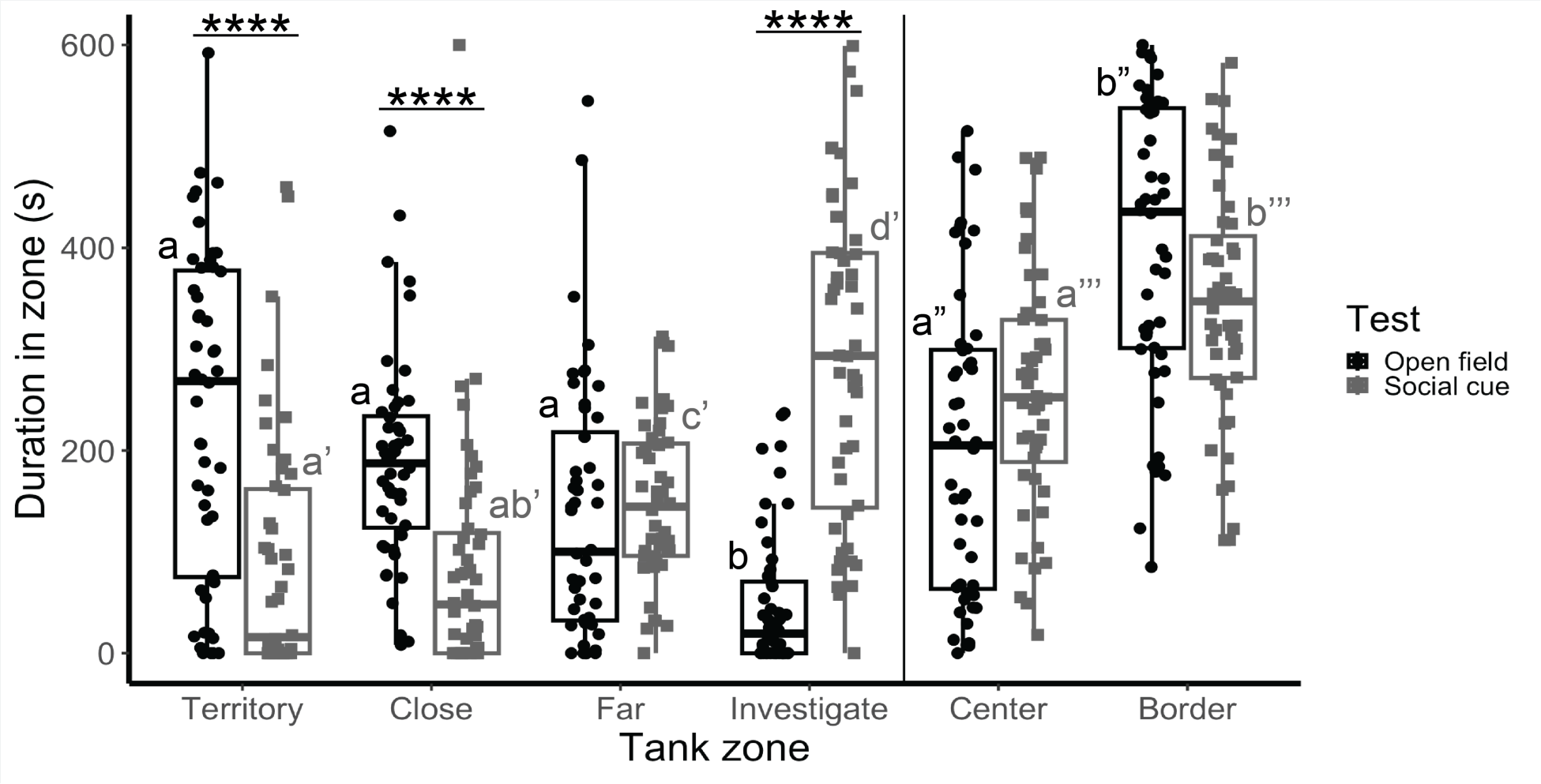
Time spent (s) in each aquarium zone, including the territory, close, far, and investigation zones of the aquarium during the open field exploration and social cue investigation. Center and border zones overlap with the other zones (see aquarium layout in Fig 1A) and were analyzed separately. Different letters indicate significant differences. Single (’), double (’’), or triple (’’’) marks indicate separate analyses. *p<0.05, **p<0.01, ***p<0.001.

**Table 1:** P-values of *post hoc*, pairwise comparisons of how long fish spent in each zone of the aquarium during the open field exploration and social cue investigation. Pairwise comparisons between aquarium zones in the open field exploration in the upper triangular (values in black). Pairwise comparisons between aquarium zones in the social cue investigation in the lower triangular (values in grey). Significant p-values are bolded.

|  |  | Territory | Close | Far | Investigation |
| --- | --- | --- | --- | --- | --- |
|  |  | Open field exploration |  |  |  |
| Territory | Social cue invest. |  | 0.59 | 0.06 | **<0.0001** |
| Close |  | 0.99 |  | 0.13 | **<0.0001** |
| Far |  | **0.005** | **0.00087** |  | **<0.0001** |
| Investigation |  | **<0.0001** | **<0.0001** | **<0.0001** |  |

We also tested for differences in how long juveniles spent in a given zone between the open field exploration and social cue investigation (Figure 3). We found that juveniles spent significantly more time in the territory zone (p<0.00001, r=0.66) and the close zone (p<0.00001, r=0.70) during the open field exploration. There was no significant difference in the time spent in the far zone between the open field and social cue tests (p=0.22). Juveniles spent significantly more time in the investigation zone, near the social cue, during the social cue investigation (p<0.00001, r=0.84). There was no significant difference in the time spent in the center between the open field exploration and social cue investigation (p=0.15). See Supplemental Information for the frequency of fish entering each aquarium zone (Supplemental Figure 1, Supplemental Table 3).

Next, we used PCA to identify behavioral patterns of where juveniles spent their time across the open field exploration and social cue investigation tests. In the analysis, we included time in each zone (territory, close, far, investigation) and time in the center for both tests. We found that the top two PCs accounted for 56.8% of the variation in the data, including PC1 (33.7%) and PC2 (23.1%). The top behavior variables that load on PC1 and PC2 are summarized in Figure 4A. Briefly, for PC1, time in the territory (open field and social cue), time in the close zone (social cue), and time in the center (open field and social cue) loaded strongly in the same direction, with time in the investigation zone (open field and social cue) and time in the far zone (social cue) loading in the opposite direction. Unlike PC1, in which space use in the open field exploration mirrored the social cue investigation, PC2 reflects a change of behavior between the tests. Time in the far zone and investigation zone in the open field exploration, and time in the territory zone and center in the social cue investigation, loaded together strongly on PC2 in the same direction. Time in the territory zone in the open field exploration and time in the investigation zone in the social cue investigation loaded strongly in the opposite direction on PC2.

**Figure 4:**
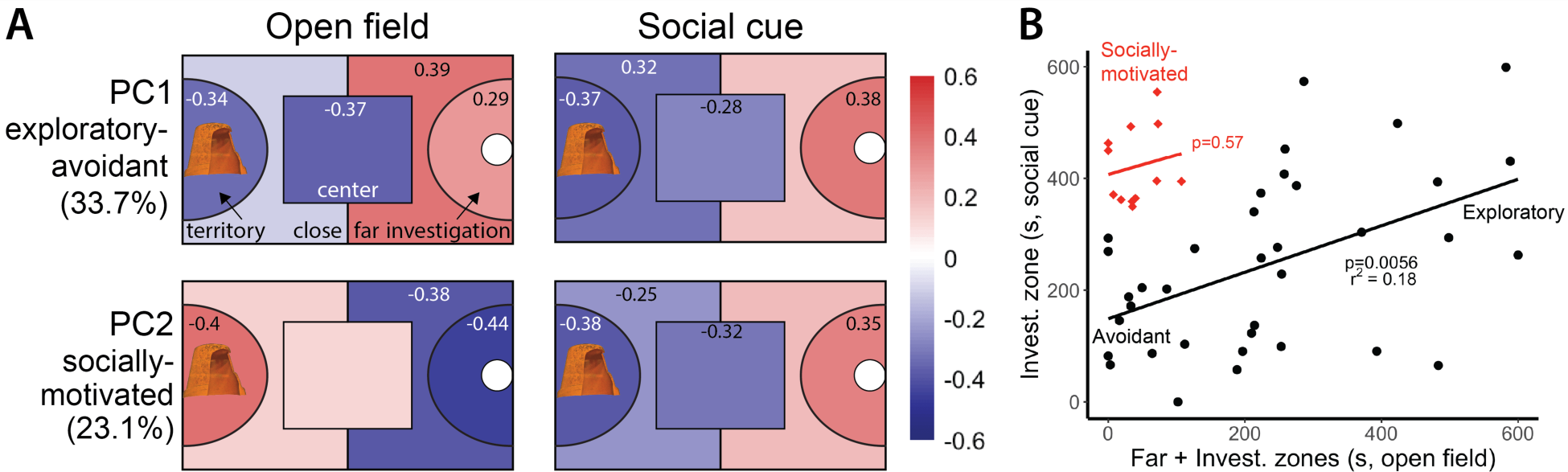
Principal components analysis (PCA) of behavior in the open field exploration (OF) and social cue investigation (SC), including time (s) in each zone (territory, close, far, investigation) and time (s) in the center for both tests. **A.** Summary of eigenvalues mapped onto the associated aquarium zones for the open field exploration and social cue investigation. Numerical values are shown for the PCA variables that load strongly on PC1 (33.7%) and PC2 (23.15) (eigenvalues stronger than ±0.25). See Figure 1.B.i. for an accurate layout of the aquarium zones. **B.** Association between time in the far + investigation zones in the open field exploration and time in the investigation zone in the social cue investigation. Fish in the 1st-3rd quartiles of PC2 are shown in black circles along an exploratory-avoidant behavioral syndrome. Fish in the 4th quartile of PC2 are shown in red diamonds (socially-motivated).

We found that different individuals drove the behavior patterns observed in PC1 vs. PC2. Fish in the first, second, and third quartiles of PC2 exhibited behavior described by PC1. For example, the PCA shows that time in the open field far and investigation zones loads in the same direction as time in the investigation zone in the social cue investigation, and these times are strongly and positively correlated for fish in the first, second, and third quartiles of PC2 (p=0.0056, r^2^=0.18). We describe the behavior type of these fish along an exploratory-avoidant continuum (Fig 4B). Fish in the fourth quartile of PC2 spend little time in the open field far and investigation zones, but in the social cue investigation, their behavior changes, spending the majority of the observation in the investigation zone (p=0.57).

### 3.3 Effects of stressors on water-borne cortisol

We found that cortisol levels differed significantly across treatment groups in response to common lab stressors (χ²=15.41, df=3, p=0.0015, η^2^=0.11, Fig 5A, Supplemental Figure 2). *Post hoc* analysis showed that cortisol levels in the net (p=0.00075) and beaker (p=0.018) treatments were significantly higher than the controls. The net treatment was also significantly higher than the orbital treatment (p=0.040). There were no significant differences between orbital and control (p=0.059), net and beaker (p=0.11), or net and orbital (p=0.42). For the beaker, net, and orbital treatments, we collected three water-borne hormone samples per fish (Fig 5B,C, Supplemental Fig 3). We found no main effect of time period (F(2,52)=0.36, p=0.71) or treatment (F(2,26)=1.59, p=0.22) on cortisol levels, and there was no interaction effect between time and treatment (F(4,52)=0.44, p=0.78).

**Figure 5:**
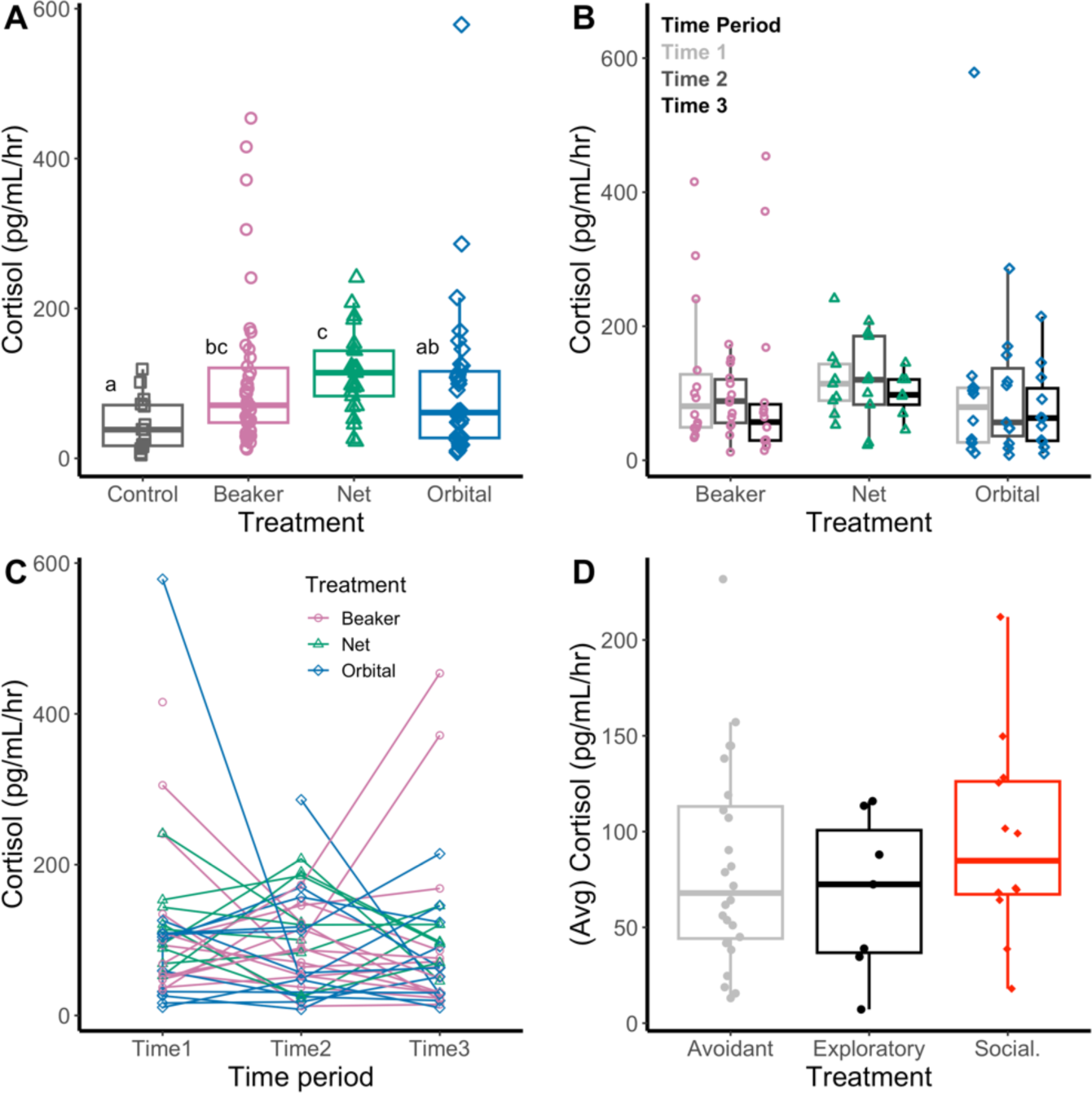
Water-borne cortisol levels (pg/mL/hr). **A.** Cortisol levels in juveniles exposed to different stressors from all time points (control: n=15 fish; beaker: n=15 fish at 3 time points; net: n=9 fish at 3 time points; orbital: n=12 fish at 3 time points). Different letters indicate significant treatment differences. **B.** Cortisol levels for juveniles in the beaker, net, and orbital treatments across hormone collection time points (time 1: 0-30 min; time 2: 30-60 min; time 3: 60-90 min). **C.** Cortisol levels for juveniles across hormone collection time points. The lines connect data points from the same individual. **D.** Cortisol levels for juveniles identified in the behavior tests as avoidant, exploratory, or socially-motivated (social.). For fish in the beaker, net, or orbital treatments, cortisol is the average of the three time points.

Because any variations in individual handling or experimental protocol can affect stress physiology, we also tested for the effects of potential confounds. Midway through the experiment, we shifted to using custom-made, small nets to transfer the fish. There were no significant differences in cortisol levels between fish handled with large versus small nets (t=-0.65, df=108.43, p=0.52). Finally, a few fish jumped out of the net and onto the lab bench when being transferred between aquaria or water-borne hormone collection beakers. There were no significant differences in cortisol levels between fish that experienced a fall at any point during the hormone collection process compared to fish that did not (t=-0.48, df=39.59, p=0.64).

### 3.4 Associations between behavior and cortisol

Finally, we tested whether behavior in the open field exploration and/or social cue investigation was associated with cortisol. To analyze the control group, which only had one hormone collection, together with the beaker, net, and orbital groups, which had three hormone collections, we calculated the average cortisol levels for the beaker, net, and orbital groups. We did not find any significant associations between cortisol with zone crosses (as a proxy for activity / locomotion) in the open field exploration or social cue investigation; time spent in the center in the open field exploration or social cue investigation; or with PC1 or PC2 (Table 2, Supplemental Figure 4). There were no significant associations between cortisol with time or frequency in any aquarium zones, or with specific cortisol collection times (1, 2, or 3) (data not shown). To test for differences by behavior type, we divided the fish along the exploratory-avoidant continuum so that exploratory fish spent >300 s (half of the observation) in the far and investigation zones in the open field test, and avoidant fish spent < 300 s. We found no significant differences in cortisol among exploratory, avoidant, or socially-motivated fish (F(2,40)=0.68, p=0.51).

**Table 2:** Associations between candidate behavior and cortisol (pg/mL/hr) for all treatment (control, net, beaker, orbital) groups together. P-values and r-squared values from Pearson correlations. Total number of zone line crosses was used as a proxy for activity / locomotion. OF: open field exploration; SC: social cue investigation; PC: principal component.

| **Cortisol x behavior** | **p-value** | **r<sup>2</sup>** |
| --- | --- | --- |
| Zone crosses (OF) | 0.11 | 0.26 |
| Zone crosses (SC) | 0.80 | -0.02 |
| Time in center (OF) | 0.71 | -0.02 |
| Time in center (SC) | 0.30 | -0.002 |
| PC1 (33.7%) | 0.49 | -0.01 |
| PC2 (23.1%) | 0.64 | -0.1 |

## 4. Discussion

We investigated early-life behavior and HPI axis function in juvenile *A. burtoni* less than 1-week old. We first demonstrated that repeated exposure to beaker confinement, which is part of the water-borne hormone collection protocol, did not lead to lower cortisol levels. Adults of other cichlid species habituation with this number of exposures (Antunes et al., 2021; Wong et al., 2008), suggesting that these young fish do not yet habituate or require more exposures and/or more time to do so. Given this outcome, we did not use the habituation protocol in the subsequent experiment. In a separate cohort of juveniles, we next demonstrated that exposure to common lab stressors (beaker and net) significantly elevated cortisol, consistent with studies in other species (Adamante et al., 2008; Cerqueira et al., 2021; Cockrem, 2013; Wong et al., 2008). Net confinement evoked the highest cortisol response. Brief movement on the orbital shaker led to the lowest cortisol levels of the experimental treatments, which did not differ from controls. This suggests that juveniles mount a stress response at this developmental stage at this young age (Botía et al., 2023; MacDougall-Shackleton et al., 2019). For most fish (∼75%), behavior was similar between the open field exploration and social cue investigation, falling along an exploratory-avoidant continuum (Conrad et al., 2011). The other ∼25% of fish were socially motivated and remained in or near the territory in the open field but were highly investigative with the social cue. Cortisol did not differ across behavior types (exploratory, avoidant, socially motivated), and individual variation in behavior was not associated with cortisol. Overall, these findings offer valuable insights into early-life stress physiology and behavior during a life history stage when experiences are known to shape development in *A. burtoni* (Harmon et al., 2024; Solomon-Lane and Hofmann, 2019) and across species (Denver, 1997; Emmerson, 2022; Gardner et al., 2009).

A persistent challenge in stress research is that the methods used can, themselves, serve as stressors (Balcombe et al., 2004; Langkilde and Shine, 2006; Romero, 2004). Here, we measured water-borne cortisol, collected from a fish’s holding water, which strongly correlates with hormone levels in blood samples (Friesen et al., 2012; Kidd et al., 2010). Although this method is non-invasive, in adult daffodil (*N. pulcher*) and convict cichlids (*A. nigrofasciata*), placement in a beaker for an extended period (30 min - 2 hrs) led to increased cortisol. However, repeated exposure to the hormone collection beaker over 3 to 4 days led to a significant decrease in cortisol, suggesting that water-borne cortisol following habituation may provide a more accurate measure of baseline levels (Antunes et al., 2021; Wong et al., 2008). We found no differences in cortisol among juveniles exposed to the water-borne hormone collection protocol 4 days in a row compared to handled and unhandled control groups. Because we only measured cortisol on day 4, rather than during each exposure (as in (Antunes et al., 2021; Wong et al., 2008), we cannot yet determine whether juveniles of this age do not habituate or if additional exposures and/or time are needed for habituation to occur. We plan to investigate this directly in future experiments. If we find that young juveniles habituate following more exposures, we would weigh the benefits of using a habituation protocol against disadvantages specific to a future experiment. For example, comparing baseline cortisol to stress-induced cortisol, or cortisol following ACTH challenge, can uncover HPA/I axis temporal dynamics (Kim et al., 2018; Lane et al., 2021; Phillips and Klukowski, 2008; Romero, 2004). However, any multi-day protocol would necessarily expose fish to repeated stress, preclude studying fish younger than the time required for habituation (e.g., 1-day old fish), and could interfere with detailed manipulations of early-life experience (e.g., (Harmon et al., 2024).

In a separate cohort of juveniles, we next tested behavior, cortisol responses to specific stressors, and the associations between behavior and cortisol levels. The young fish readily engaged in both the open field exploration and social cue investigation and showed location preferences similar to older *A. burtoni* juveniles (Harmon et al., 2024). In the open field, fish spent the most time in the territory zone, and duration decreased with distance from the territory. Focal fish altered their behavior when the social cue was introduced into the aquarium, spending significantly more time near the novel juvenile. The strong salience of a social signal for young juvenile *A. burtoni* is consistent with this species being highly social across life history stages (Fernald and Hirata, 1979; Fraley and Fernald, 1982; Maruska and Fernald, 2018). Principal components analysis revealed interesting patterns of where fish spent time in the aquarium. Principal component 1 (33.7%) showed similar behavior between the open field and social cue investigation tests. Fish that spent more time in the far and/or investigation zones in the open field also spent more time in those zones, and away from the territory, with the addition of the social cue. These data align with a behavioral syndrome we previously identified in older juvenile *A. burtoni*. That syndrome also included dominance behavior, which was not measured in this study (Harmon et al., 2024; Solomon-Lane and Hofmann, 2019).

Principal component 2 (23.1%) revealed a behavior pattern we have not previously identified. A subset of individuals (∼25%) remained in the territory and close zones for the open field exploration but then moved closer to the social cue (far and investigation zones) for the duration of that observation. Open field behavior has been interpreted as exploration-avoidance or boldness-shyness behavioral syndromes, depending on how the test was carried out, the study species, and experimental context (Carter et al., 2013; Conrad et al., 2011; Dall et al., 2012; Réale et al., 2007; Toms et al., 2010; Walsh and Cummins, 1976). In identifying these two behavior types, we confirm that these tests are measuring distinct features of behavioral phenotype. Given our testing conditions, which placed fish directly in the open field without the choice of whether to enter, and the absence of any risk (e.g., predation signal), we suggest this behavior is more likely exploration-avoidance, rather than boldness-shyness. However, further validations are needed to contextualize our findings (Carter et al., 2013; Réale et al., 2007).

We found that the beaker and net treatments led to significant increases in cortisol levels, suggesting that these experiences were stressors for *A. burtoni* under 1 week of age. While beakers are regularly used in water-borne hormone collection protocols (Antunes et al., 2021; Wong et al., 2008), we did not know whether the handling and confinement would induce stress in these young fish. Hand nets are often used as a method of confinement stress and to move fish within the lab. We found the highest average cortisol in the net treatment. Across species, net confinement (in and out of water) is a consistently intense stressor (Zuberi et al., 2014). Cortisol following the orbital shaker was not significantly different from controls. Interestingly, water movement has context-dependent effects. Dorado fingerlings (*Salminus brasiliensis*) placed in plastic bags on an orbital shaker exhibit stress responses (Adamante et al., 2008), while in rainbow trout, water movement acted as an enrichment factor (Villarroel et al., 2021). Although many studies across species demonstrate an increase in cortisol following acute stressor exposure, these results are valuable for multiple reasons. First, they demonstrate HPI axis functionality in fish at this age. Some species experience hyporesponsive phases, such as juvenile rainbow trout (Barry et al., 1995). As a mouthbrooding species that breeds readily in the laboratory, *A. burtoni* are accessible at all larval stages, and we plan to compare these findings to HPI axis function at earlier developmental timepoints. Second, these data allow us to establish predictable ranges of cortisol. The overlap in cortisol levels between the stressor treatments with the control group suggests the beaker and net treatments were mild or moderate stressors. The stressor intensity could potentially be increased, for example, by confining a fish in a net briefly held out of the water. We also observed that these juveniles had higher cortisol levels than adults or older juvenile *A. burtoni*. Given cortisol’s role in growth and development, we hypothesize that these cortisol levels are serving a developmental function. Third, we chose these potential stressors because of their regular use in the lab, and our findings provide context for how they can affect HPI axis function. Effects of an experimental stressor could compound or interact with any background HPI axis activation from such experiences (e.g., transfer in a hand net). Finally, this foundational knowledge of HPI axis function will allow us to investigate early-life plasticity, for example, in response to predictable and unpredictable stress regimens (Galhardo et al., 2011).

We were unable to quantify the baseline, peak, and recovery phases of the stress response by collecting three sequential water samples. The handling involved in transferring fish between the collection beakers may have prolonged or re-activated the stress response, or delayed negative feedback. This is consistent with *N. pulcher,* and those researchers developed a method for draining water samples and refilling the holding container without moving the fish (Antunes et al., 2021). We plan to try this approach in the future. Negative feedback efficiency impacts how much cumulative cortisol individuals are exposed to over time and can be affected by early-life experiences (Champagne and Curley, 2005; Francis et al., 1999). In the control group, we expect that negative feedback mechanisms started during the 90-min collection period. Both control and beaker fish were treated identically in the first 30 min, after which the beaker fish were transferred for their second collection. Once corrected to be in the same units (pg cortisol / mL water in the collection beaker / hr collection time), control fish cortisol was significantly lower than for beaker fish, suggesting cortisol production decreased over the 90 min. Beaker fish cortisol was lower in the third collection, while cortisol across the timepoints for net and orbital fish were similar. There were no significant effects of time. Comparing individual cortisol levels across collection timepoints for the beaker, net, and orbital fish showed striking differences in patterns of cortisol changes. Some fish exhibited consistent levels across time, while others showed a “V” or inverted ”V” pattern. The patterns do not appear specific to stressor-type. In future studies, we aim to uncover the factors shaping these differences in stress response, such as early-life experiences and/or genetics (Cockrem, 2013). Here, incidental experiences, such as a fall between collection beakers, did not explain this variation.

We next assessed whether behavior in the open field exploration and/or social cue investigation tests prior to stress exposure was associated with individual variation in cortisol. Interestingly, we found no associations between behavior and cortisol levels, including with zone crosses as a proxy for locomotion / activity, time in the center, or behavior PCs. The relationship between cortisol and behavior is context-specific; therefore, results vary across studies. Locomotion and/or activity are often described as anxiety-like behaviors that are positively associated with glucocorticoids under some conditions. In Wistar rats (*Rattus norvegicus*) (Sandi et al., 1996), white-crowned sparrows (*Zonotrichia leucophrys gambelii*) (Breuner et al., 1998), and zebrafish (*Danio rerio*) (Ghisleni et al., 2012), locomotion was positively associated with glucocorticoid levels. In rainbow trout, short-term exogenous cortisol treatment increased locomotion in juveniles challenged by a conspecific intruder, but had no effect on the locomotion of socially undisturbed fish (Øverli et al., 2002). In our study, zones were drawn around the territory and social cue placement and may not be well-suited as a proxy for locomotion / activity. Time spent in the center of the aquarium is an indicator for multiple behavior states. Exploratory and/or bold individuals can spend more time in the center, often associated with lower levels of HPA/I axis activation. Anxiety-like or fear-related behaviors might result in either increased time spent near the outer edge or increased time in the center as a consequence of increased locomotion, which is often associated with higher HPA/I axis activation. We expected to find lower cortisol levels in the exploratory fish and higher cortisol levels in the avoidant fish; however, we did not find a difference. For the socially-motivated fish, we do not yet know if lack of exploration in the open field was due to anxiety/fear or lack of incentive.

In identifying multiple behavior types in such young fish, we have the opportunity to test for phenotypic differences that persist long-term. For example, the exploratory juveniles could be proactive copers. Coping styles refer to a suite of behaviors forming a syndrome that aligns with stress physiology (Koolhaas et al., 2010, 1999). In adult *A. burtoni*, individuals switch between reactive and proactive behaviors, rather than remaining consistent across contexts (Butler et al., 2018). The juveniles could also be experiencing a fast “pace-of-life,” which can include behaviors, such as increased activity, exploration, boldness, and aggressiveness, as well as traits that are part of the slow-fast metabolic continuum (Careau and Garland, 2012).

## 5. Conclusion

In the present study, we asked fundamental questions about HPI axis function in very young juvenile *A. burtoni*. Previous work has demonstrated early-life social effects on HPI axis function and associations with behavior (Harmon et al., 2024; Solomon-Lane and Hofmann, 2019); however, there were critical gaps in our knowledge. Here, we show that juveniles respond to the acute stress of net and beaker confinement with an elevation in cortisol. In measuring cortisol in response to common lab experiences and stressors, we can use these data in interpreting future results, particularly in disentangling the effects of experimental manipulations from a stress response. Juveniles that were repeatedly exposed to beaker confinement, a necessary component of the water-borne hormone collection method, did not habituate (or sensitize) to the experience. Future studies can build on our findings to ask whether habituation occurs in older juveniles or following more exposures. Surprisingly, we identified two distinct behavior types, an exploratory-avoidant syndrome and a group of socially-motivated fish that changed their behavior in the presence of the social cue. This shows striking individual variation in behavior at a very young age, and we aim to investigate if behavior type is consistent through time and across contexts. Neither behavior, nor behavior type, was associated with cortisol levels. Cortisol-behavior interactions are highly context-specific, and measuring behavior pre- and post-stressor, and baseline compared to stress-induced cortisol, can further elucidate the complex relationships between hormones behavior.

## Acknowledgements

This work was funded by NSF grant IOS-2341006 to TKSL, start-up funds from the W.M. Keck Science Department to TKSL, W.M. Keck Science Department Summer Fellowship to APA, and Claremont McKenna College Sponsored Internships & Experiences to JL. We thank Veronica Britton, Samuel Chan, Cristi Cruz, Jessica Gonzalez Rodriguez, Alex Kasiske, Blake Migden, Emma Thompson, and Andrew Yuan for contributing to hormone sample collection and analysis. We thank the Solomon-Lane Lab for contributing to fish care and maintenance.

## Data availability

All data will be made public on Dryad upon publication. Private reviewer link: http://datadryad.org/stash/share/9vrdN1T_pevPtokrxy3RFM8VxLMnbRA9SMIgJ_WCNS0

