## Supplemental Information for "Early-Life Behavior Phenotypes and Cortisol Responses to Common Lab Stressors in a Cichlid Fish"

**Supplemental Table 1:** Mass (g) of experimental juveniles in the habituation experiment.

| **Treatment** | **Mean mass** | **SEM** | **Median mass** | **Min mass** | **Max mass** |
| --- | --- | --- | --- | --- | --- |
| All | 0.016 | 0.00058 | 0.016 | 0.008 | 0.023 |
| Unhandled | 0.017 | 0.00098 | 0.017 | 0.01 | 0.023 |
| Handled | 0.015 | 0.00089 | 0.015 | 0.01 | 0.021 |
| Habituated | 0.016 | 0.0011 | 0.015 | 0.008 | 0.022 |

All juveniles were 4 days old on the day of water-borne hormone collection (counting from the time fish were removed from the mother’s buccal cavity).

**Supplemental Table 2:** Mass (g) of experimental juveniles in the behavior and cortisol response to stressors.

| **Treatment** | **Mean mass** | **SEM** | **Median mass** | **Min mass** | **Max mass** |
| --- | --- | --- | --- | --- | --- |
| All | 0.019 | 0.0012 | 0.017 | 0.011 | 0.038 |
| Control | 0.02 | 0.034 | 0.017 | 0.011 | 0.038 |
| Beaker | 0.018 | 0.0018 | 0.015 | 0.012 | 0.028 |
| Net | 0.02 | 0.0024 | 0.018 | 0.013 | 0.034 |
| Orbital | 0.018 | 0.0015 | 0.018 | 0.012 | 0.024 |

All juveniles were less than 7 days old on the experimental day (counting from the time fish were removed from the mother’s buccal cavity). Juveniles first underwent the open field exploration and social cue investigation behavioral tests. They were then randomly assigned to a stress treatment for water-borne hormone collection: control, beaker, net, or orbital. Means and standard error of the mean (SEM), median, minimum (min), and maximum (max).

Juvenile behavior in the open field exploration and social cue investigation behavioral tests was observed before fish were exposed to a lab stressor and water-borne hormone sample collection. In both the open field exploration (χ²(3)= 57.5, p<0.0001, W=0.40) and social cue investigation (χ²(3)= 56.4, p<0.0001, W= 0.39), juveniles spent significantly different amounts of time in the different zones of the tank (Supplemental Figure 1). In the open field exploration, juveniles entered the close zone the most times and the investigate zone the fewest times. In the social cue investigation, juveniles entered the far zone the most times and the territory zone the fewest. Supplemental Table 3 shows the results of *post hoc* pairwise comparisons between zones for both behavior tests.

We also tested for differences in how many times juveniles entered a given zone between the open field exploration and social cue investigation. We found that juveniles entered the territory zone (p<0.0001, r=0.68) and the close zone (p<0.0001, r=0.69) significantly more times in the open field exploration compared to the social cue investigation. Juveniles entered the far zone (p=0.00041, r=0.50) and the investigate zone (p<0.0001, r=0.69) significantly more frequently in the social cue investigation compared to the open field exploration. There was no significant difference in the frequency of entering the center zone between the open field exploration and social cue investigation (p=0.054).

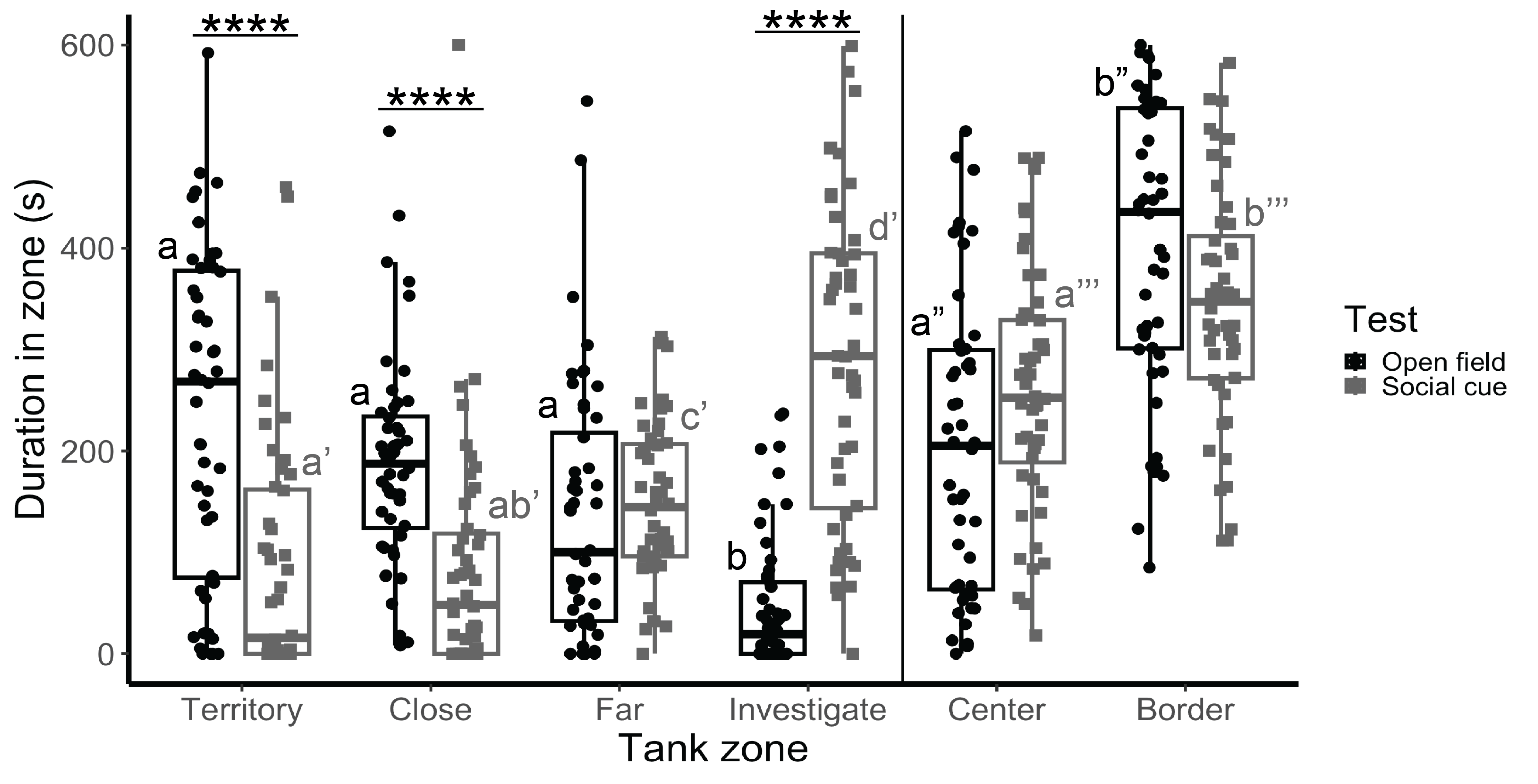

**Supplemental Figure 1:** Frequency entering each tank zone, including the territory, close, far, and investigate zones of the tank during the open field exploration and social cue investigation. Center and border zones overlap with the other zones (Fig 1A) and were analyzed separately. Different letters indicate significant differences. Single (’), double (’’), or triple (’’’) marks indicate separate analyses. *p<0.05, **p<0.01, ***p<0.001.

**Supplemental Table 3:** P-values of *post hoc*, pairwise comparisons of how frequently fish entered each zone of the tank during the open field exploration and social cue investigation.

|  |  | **Territory** | **Close** | **Far** | **Investigation** |
| --- | --- | --- | --- | --- | --- |
|  |  | **Open field exploration** | | | |
| **Territory** | **Social cue invest.** |  | **< 0.0001** | 0.76 | **0.00022** |
| **Close** |  | **<0.0001** |  | **0.0003** | **< 0.0001** |
| **Far** |  | **<0.0001** | **0.00053** |  | **< 0.0001** |
| **Investigation** |  | **0.00058** | 0.51 | **<0.0001** |  |

P-values of *post hoc* pairwise comparisons between tank zones in the open field exploration (upper triangular, values in black). P-values of *post hoc* pairwise comparisons between tank zones in the social cue investigation (lower triangular, values in grey). Significant p-values are bolded.

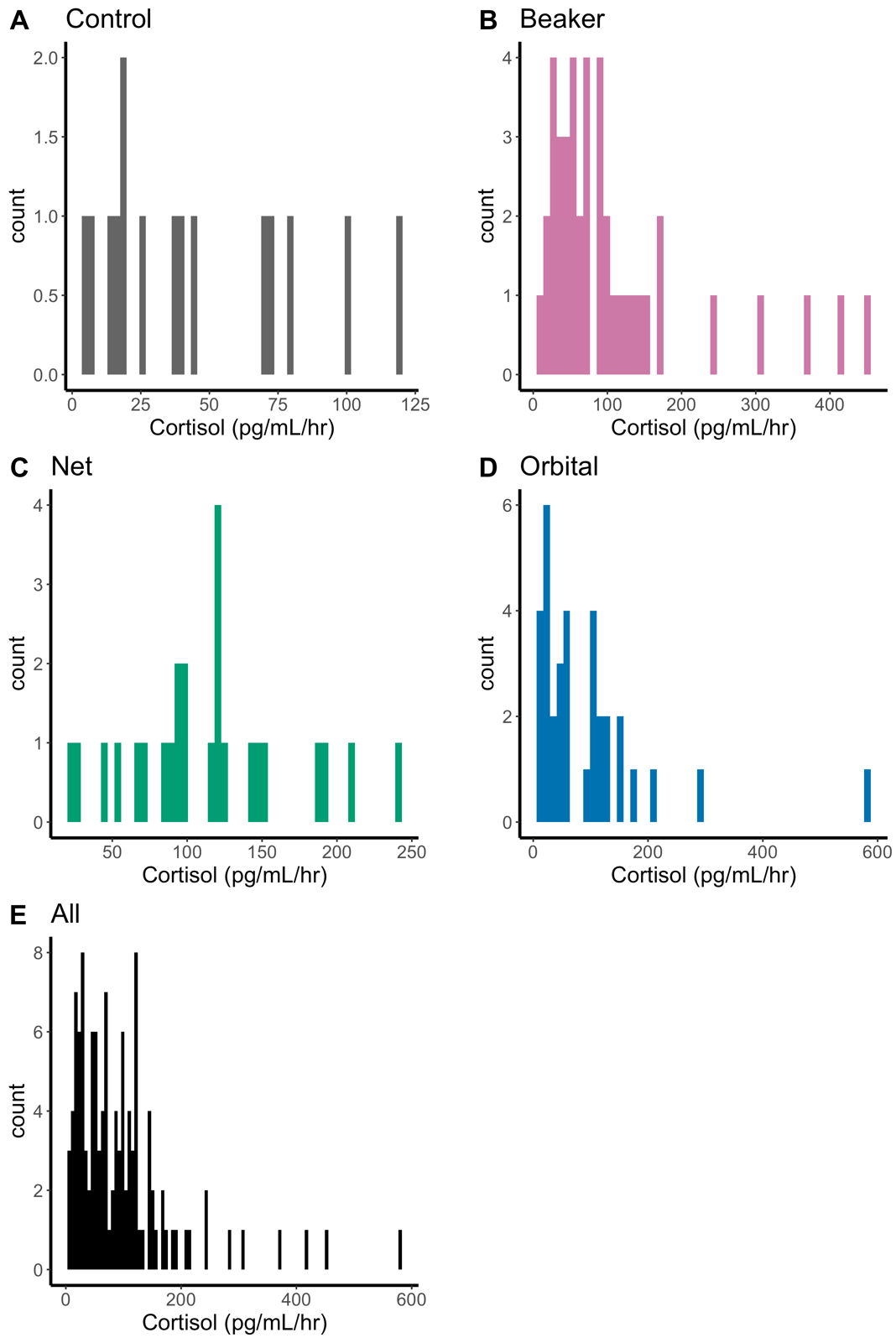

**Supplemental Figure 2:** Frequency distributions of water-borne cortisol levels (pg/mL/hr) in juveniles exposed to A) control, B) beaker, C) net, and D) orbital stressors. E) Juveniles from all treatment groups.

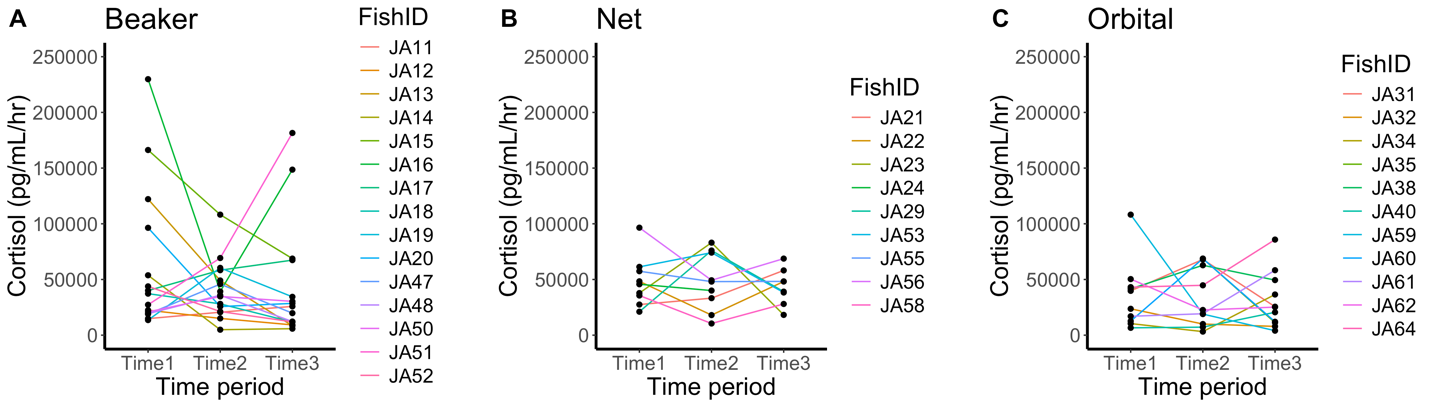

**Supplemental Figure 3:** Individual variation in water-borne cortisol across collection time points (time 1: 0-30 min; time 2: 30-60 min; time 3: 60-90 min) for fish in the A) beaker (n=15), B) net (n=9), and C) orbital (n=12) treatments.

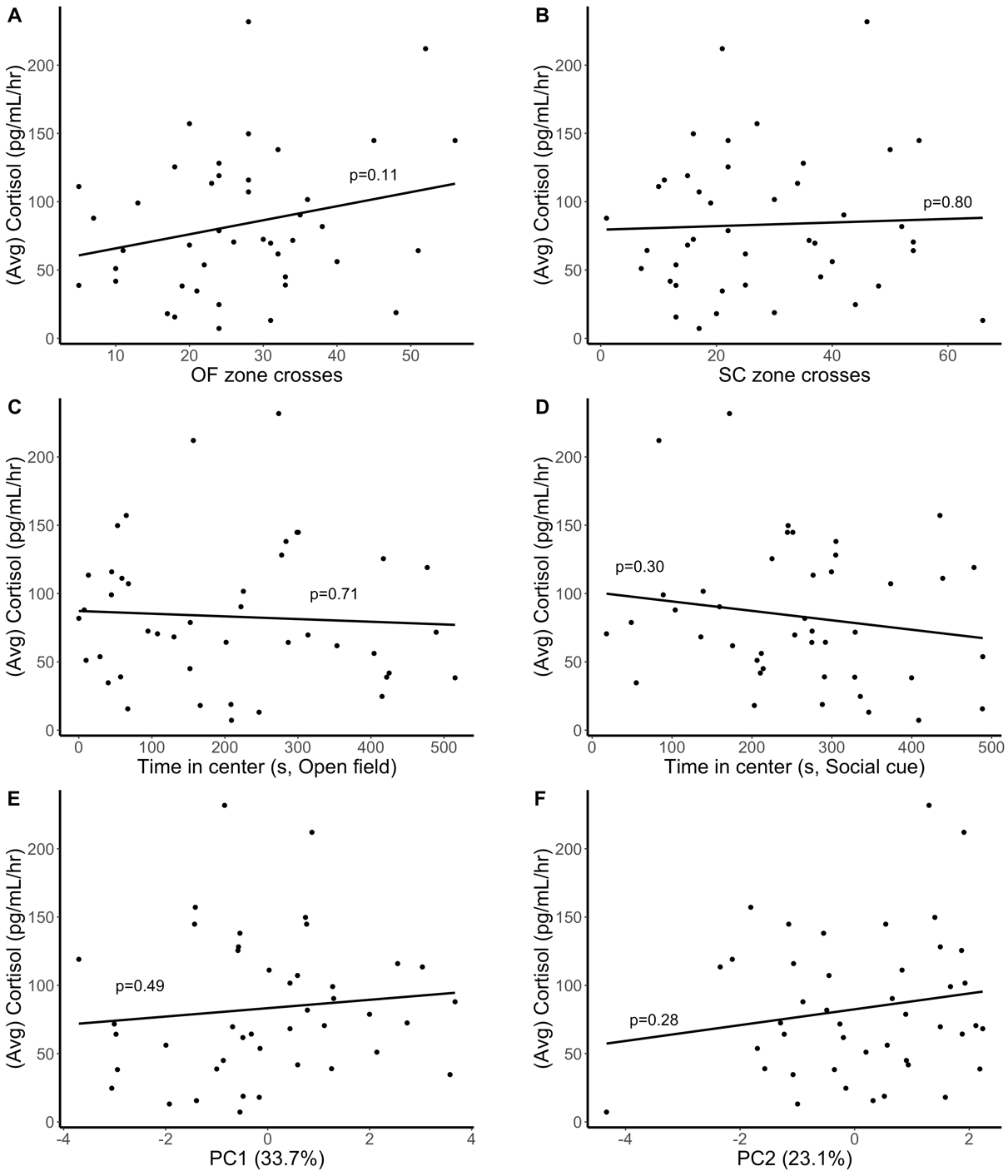

**Supplemental Figure 4:** Associations between cortisol and candidate behaviors: **A.** zone crosses in the open field (OF) exploration, **B.** zone crosses in the social cue (SC) investigation, **C.** time in the center in the open field exploration, **D.** time in the center in the social cue investigation, and with E) principal component (PC) 1 and F) PC2. The percentage of variation in the data explained by each PC is in parentheses.
